# Clonal assessment of functional mutations in cancer based on a genotype-aware method for clonal reconstruction

**DOI:** 10.1101/054346

**Authors:** Paul Deveau, Leo Colmet Daage, Derek Oldridge, Virginie Bernard, Angela Bellini, Mathieu Chicard, Nathalie Clement, Eve Lapouble, Valérie Combaret, Anne Boland, Vincent Meyer, Jean-François Deleuze, Isabelle Janoueix-Lerosey, Emmanuel Barillot, Olivier Delattre, John Maris, Gudrun Schleiermacher, Valentina Boeva

## Abstract

In cancer, clonal evolution is characterized based on single nucleotide variants and copy number alterations. Nonetheless, previous methods failed to combine information from both sources to accurately reconstruct clonal populations in a given tumor sample or in a set of tumor samples coming from the same patient. Moreover, previous methods accepted as input all variants predicted by variant-callers, regardless of differences in dispersion of variant allele frequencies (VAFs) due to uneven depth of coverage and possible presence of strand bias, prohibiting accurate inference of clonal architecture. We present a general framework for assignment of functional mutations to specific cancer clones, which is based on distinction between passenger variants with expected low dispersion of VAF versus putative functional variants, which may not be used for the reconstruction of cancer clonal architecture but can be assigned to inferred clones at the final stage. The key element of our framework is QuantumClone, a method to cluster variants into clones, which we have thoroughly tested on simulated data. QuantumClone takes into account VAFs and genotypes of corresponding regions together with information about normal cell contamination. We applied our framework to whole genome sequencing data for 19 neuroblastoma trios each including constitutional, diagnosis and relapse samples. We discovered specific pathways recurrently altered by deleterious mutations in different clonal populations. Some such pathways were previously reported (e.g., MAPK and neuritogenesis) while some were novel (e.g., epithelial-mesenchymal transition, cell survival and DNA repair). Most pathways and their modules had more mutations at relapse compared to diagnosis.

## Introduction

The principal cause of cancer is believed to be accumulation of mutations and structural variations (SVs) of the genome. Recently, many efforts have focused on the identification of driver mutations; nonetheless, passenger variants, although they are not directly linked to the disease, may provide additional evidence from which to infer the phylogeny of a tumor and so help uncover the basis for its proliferative activity (Marusyk et al., 2014).

To understand the role driver mutations play in clonal expansion and cancer progression, it is essential to accurately reconstruct the clonal structure and assign functional variants to it. We define a clone as a cell population that harbors a unique pattern of mutations and SVs. Clones are related to each other and share a common ancestor. A hierarchical phylogenetic tree, which represents the ancestry of clones, can be constructed to reflect the order of appearance of new sets of mutations defining each clone. Each such set of mutations is expected to contain at least one driver mutation or SV giving a selective advantage to the clone compared to its ancestry. A clone can thus have a different behavior from its ancestral clone when facing the same stimuli. With accumulation of driver mutations, clones are likely to gain hallmarks of cancer such as evading growth suppressors, activating invasion and metastasis (Hanahan and Weinberg, 2011).

High-Throughput Sequencing (HTS) of bulk tumor tissues has allowed uncovering genetic differences at the clonal level in primary and relapse/metastatic tumors. Modern computational methods provide ways to reconstruct the structure of the phylogenetic tree from variant allele frequencies (VAFs) in sequenced reads, where VAF is a proportion of reads supporting each given variant among all reads spanning the position of interest (Fischer et al., 2014; Jiao et al., 2014; Kepler, 2013; Malikic et al., 2015; Miller et al., 2014; Qiao et al., 2014; Schwarz et al., 2014). However, existing methods for clonal reconstruction often neglect information about the genotype of each position, which refers to the paternal or maternal inheritance of a locus and the number of copies of each allele. Accounting for the genotype information is especially crucial in the case of hyper-diploid cancers and cancers with highly rearranged genomes, as the cellular prevalence – measured as the proportion of cancer cells carrying a variant – is linked to VAF through such parameters as copy number of the locus and the number of chromosome bearing the mutation.

Here we show that by combining the genotype and VAF information it is possible to correctly cluster variants and assign them to specific clones, thus reconstructing the clonal architecture of an individual cancer. This may be done with our novel method, QuantumClone, designed to reconstruct clones based on both VAF and genotype information. We demonstrate that our algorithm accurately clusters variants on simulated data, even when cancer is hyper-diploid or contaminated by normal cells. We also propose a general framework based on QuantumClone to detect driver mutations of clonal evolution. This general approach is applied to 19 neuroblastoma cases; each case includes whole genome sequencing (WGS) data from a sample at diagnosis and relapse. We show that deleterious mutations in neuroblastoma accumulate at relapse in specific pathways such as cell motility (e.g., cell-matrix adhesion and regulation of epithelial-mesenchymal transition, EMT) and cell survival (e.g., PI3K/AKT/mTOR, MAPK or noncanonical Wnt pathways).

## Results

The QuantumClone method presented here applies an expectation-maximization (EM) algorithm and allows for accurate inference of clonal structure using VAFs from one or several tumor samples sequenced using WGS. It can analyze variants coming from highly rearranged and hyper-diploid cancer genomes. We extensively validated QuantumClone on simulated data, where we compared it with recently published methods (Miller et al., 2014; Roth et al., 2014). We complement QuantumClone with a robust framework for the functional assessment of mutations based on signaling pathway analysis combined with the clonal assignment (Fig. 1).

**Figure 1:**
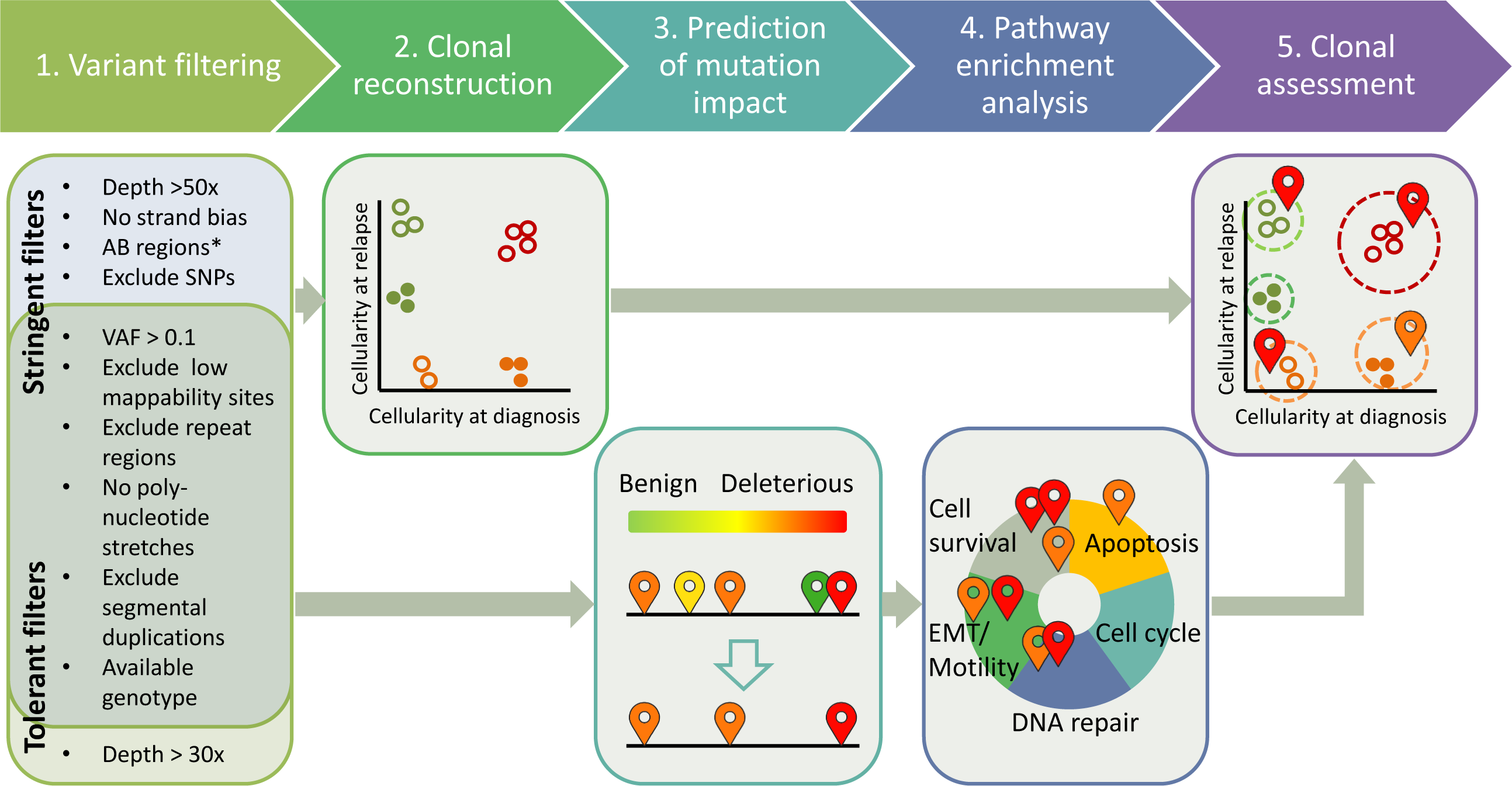
Overview of the general clonal reconstruction workflow: steps 1-5. (1) Variants arc filtered to remove false positive calls; stringent filters are used to produce mutations that are further employed for clonal reconstruction (step 2). tolerant filters are used to detect functional mutations (step 3-4). (2) Variants that pass stringent filters and have genotype information assigned to the corresponding genomic loci are used as input to QuantumClone to reconstruct clonal populations. (3) Functional impact of variants passing tolerant filters is assessed. (4) Pathways recurrently altered by deleterious mutations are identified. (5) Finally, possibly damaging mutations belonging to frequently altered pathways are mapped to the reconstructed clones. (*) Stringent filtering keeps mutations located in AB regions only when at least 100 mutations pass this filter.

The overall framework was applied to WGS neuroblastoma datasets: 19 patients’ primary and relapse samples including 7 new triplets. Novel and previously published samples (Eleveld et al., 2015) have been sequenced at ~ 100 × depth of coverage using Illumina HiSeq 2500 and Complete Genomics sequencing technologies. Application of the QuantumClone-based framework allowed us to discover pathways recurrently altered by mutations in neuroblastoma at diagnosis and relapse.

### Assessment of clonal reconstruction accuracy by QuantumClone

For clonal reconstruction using VAFs, we developed an approach that applies an EM algorithm (Fig. 2A, Methods). QuantumClone utilizes genotype information and assigns variants to clones providing the most likely values of cellular prevalence (Fig. 2A, Methods).

**Figure 2:**
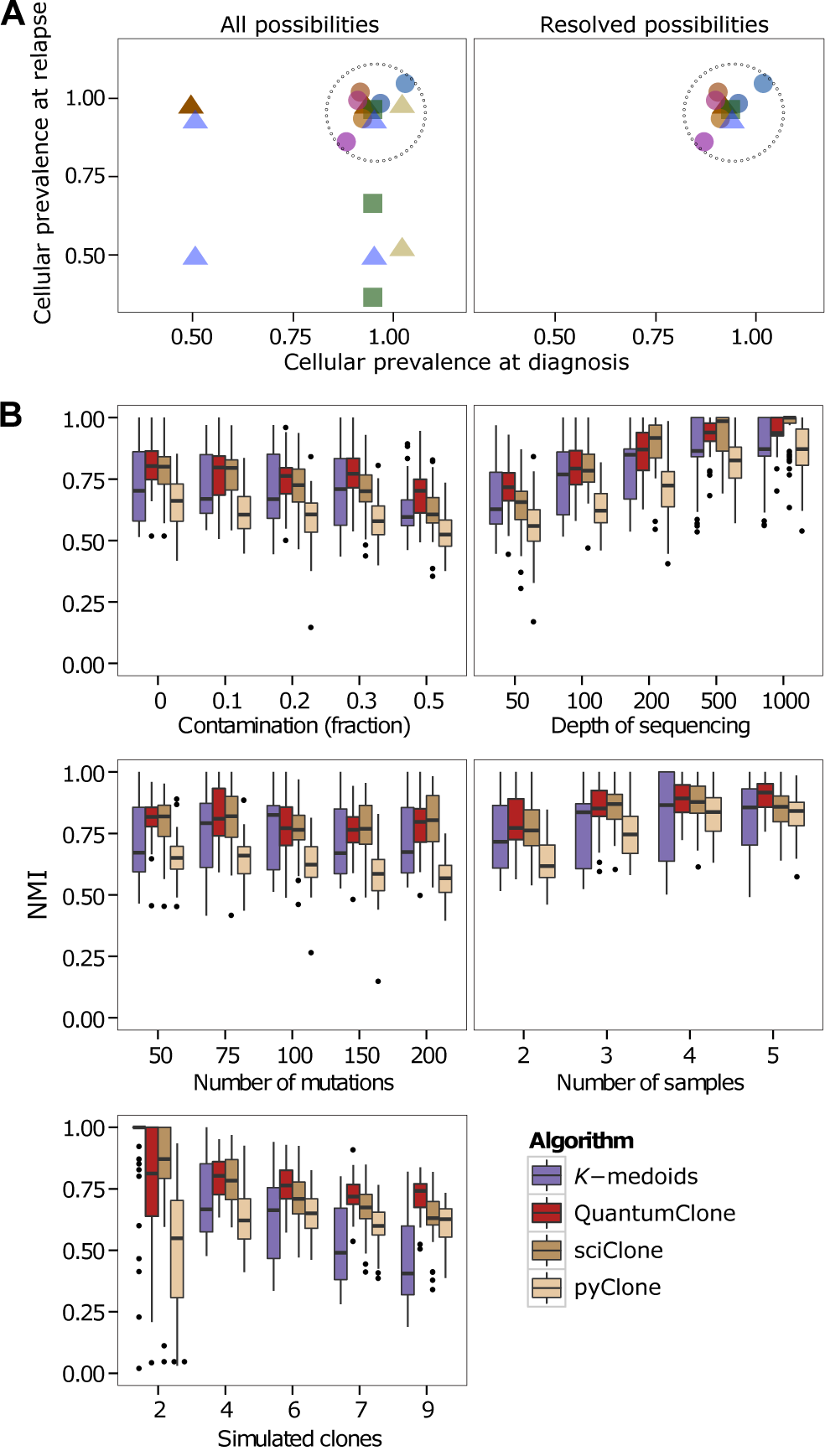
Principle of the QuantumClone algorithm and comparison to published methods. **(A)** Mutations located in regions of copy number aberration can be present on several chromosomal copies: they can thus be assigned to several cellular prevalence values (left panel). After the Expectation Maximization (EM) step each mutation is attributed the most likely cellular prevalence value (right panel). Each mutation is represented by a specific color. Mutations located in AB regions (circles): mutations located at relapse in regions of gain (squares), mutations located in regions of gain both at diagnosis and relapse (triangles). **(B)** Comparison of QuantumClone to existing methods. Normalized Mutual Information (NMI) is used to assess the quality of clustering on simulated data, with a single parameter varying in each test. QuantumClone (red) shows better performance in difficult settings, i.e. in presence of a high number of clones, low number of input samples, low sequencing depth, and high fraction of contamination by normal cells. Default parameters: two tumor samples without contamination sequenced at 100×; 4 clones; 100 mutations used for clustering.

### Comparison of QuantumClone with existing methods

Using *in silico* data, we compared the performance of QuantumClone, sciClone (Miller et al., 2014), pyClone (Roth et al., 2014) and a generic *k*-medoids clustering algorithm in inferring clonal structure of a set of tumors derived from the same patient. sciClone is based on variational Bayesian Mixture Models, while pyClone relies on a hierarchical Bayes statistical model. Partitioning with *k*-medoids is a more robust version of a widely used *k*;-means clustering algorithm (Kaufman and Rousseeuw, 1987).

In our simulation experiments, the following parameters were varied within realistic ranges: depth of sequencing (50 × to 1000×), fraction of contamination by normal cells (from 0 to 50%), number of variants used for the clonal reconstruction (from 50 to 200), number of tumor samples used for each patient (from 2 to 5) and number of distinct clones per cancer (from 2 to 9) (Fig. 2B). For each set of parameters, we performed and analyzed 50 independent simulation experiments (Methods). The accuracy of clonal reconstruction was assessed by evaluation of the normalized mutual information (NMI) (Manning et al., 2008). A perfect mutation clustering would result in a NMI value of 1, which corresponds to an identification of the exact number of clones and correct assignment of all the mutations of a clone to the same cluster.

Our analysis showed that QuantumClone surpasses both published algorithms in experiments with challenging parameter settings: high contamination by normal cells, moderate depth of sequencing or high tumor heterogeneity (Fig. 2B). Indeed, in samples with 50% contamination by normal cells QuantumClone significantly outperformed sciClone and pyClone (*p* — *value* = 2.6 × 10^−4^ and *p* — *value* = 3.2 × 10^−14^, Welch two sample t-test). While at high values of sequencing depth, all methods provided accurate results, at depth of sequencing of 50 × QuantumClone consistently gave better predictions (*p* — *value* = 1.2 × 10^−3^ and *p* — *value = 7.3 ×* 10^−10^ for sciClone and pyClone respectively). In addition, compared to the other methods, QuantumClone took the best advantage of data when multiple samples were provided for the analysis (*p* — *value* = 9.4 × 10^−3^ mid *p* — *value* = 8.0 × 10^−4^ for sciClone and pyClone respectively, for simulated tumors with five samples). Also, starting from six clones per tumor, QuantumClone demonstrated significantly better clonal reconstitution accuracy than the other methods (*p* — *value* = 4.3 × 10^−8^ mid *p* — *value* < 2.2 × 10^−16^ for sciClone and pyClone respectively).

### Assessment of clonal reconstruction accuracy in hyper-diploid cancers or cancers with highly rearranged genomes

We expect that in addition to the parameters discussed above, the degree of genome rearrangement and chromosome duplication significantly affects the quality of the mutation clustering and consecutive clonal reconstruction. Indeed, given an observed VAF value, a mutation occurring in a high copy number locus has more possibilities for values of cellular prevalence: a mutation with an observed allele frequency of 25% can only be linked to a cellular prevalence of 50% in a AB locus, while it can arise from cellular prevalence values of 33.3%, 50% or 100% if the genotype is AAAB (Methods).

In order to validate QuantumClone on diploid and hyper-diploid genomes, we simulated variants in loci of genotype AB, AAB and A ABB (Fig. 3). In addition to QuantumClone, we tested the performance of the *k*-medoids clustering algorithm, and two alternative versions of QuantumClone: QuantumClone-Single and QuantumClone-Alpha. QuantumClone-Single assigned variants to a single copy state, e.g., variants in AAB regions were supposed to only occur on a single chromosome. QuantumClone-Alpha used the same EM algorithm as the default version of QuantumClone but added additional weights to probabilities based on the locus genotype (Methods); e.g., this method suggested that a mutation in an AAB region has 3 times more chances to occur on a single chromosome than on two chromosomes out of three. The default version of QuantumClone assigned equal weights to all possibilities (Methods).

**Figure 3:**
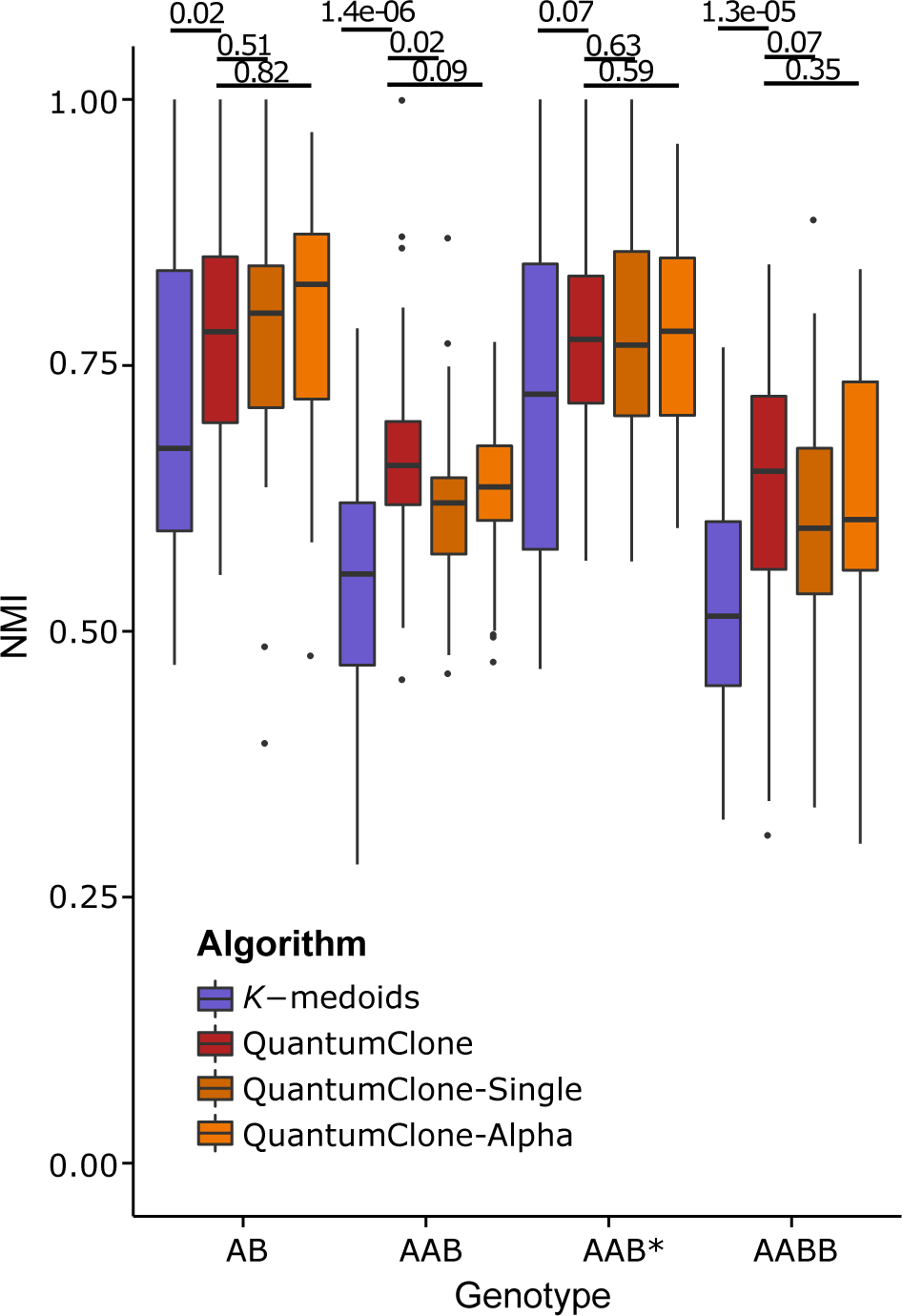
Quality of clonal reconstruction for mutations located in regions of altered copy number. QuantumClone-Single assumes that the mutation is always present at a single chromosomal copy: QuantumClone-Alpha uses an alternative algorithm for the selection of the best cellular prevalence given VAF values (Methods). Our comparison shows that the default version of QuantumClone performs as good as QuantumClone-Single in single copy regions but is the best algorithm in the situation when there are multiple possibilities of cellular prevalence for a mutation. P-values are calculated using Welch’s t-test. (*) Mutations in AAB* regions are always present at a single copy.

In all types of regions, QuantumClone and its alternative methods performed better than the baseline *k*-medoids clustering algorithm (Fig. 3). For AB regions, where VAF of each mutation corresponds to a single possible value of cellular prevalence, the difference in performance between QuantumClone, QuantumClone-Single and QuantumClone-Alpha was not significant; in fact, it was due to random initialization of the EM algorithm.

In the case of a triploid genome when each mutation was simulated to occur in a single chromosome copy (case AAB*, Fig. 3), QuantumClone provided equally good results as QuantumClone-Single (t-test *p*— *value* = 0.63 with a mutation.

We demonstrated that in a more realistic case when a mutation can have a single or multiple copy status (AAB and A ABB regions), QuantumClone performed better than the three other methods. This validated our computational strategy for hyper-diploid cancers.

Overall, validation on simulated data showed that (1) QuantumClone can be applied to cancer samples with hyperploid or rearranged genomes and (2) QuantumClone performs generally better than its peers in difficult settings, e.g., in experiments with low depth of sequencing, when the number of clones is higher than or equal to six, or when the contamination by normal cells is higher than or equal to 0.3.

### Effect of experimental settings on the clonal reconstruction accuracy

Our analysis allowed us not only to compare QuantumClone to published methods and define the limits of applicability of each method but also to study the effect of experimental settings on the clonal reconstruction accuracy, which can help in the planning of tumor DNA sequencing experiments.

#### Effect of contamination by normal cells

As expected, the accuracy of clonal reconstruction decreased with the contamination level (Spearman’s rank correlation rho = −0.29, *p — value* = 8.1 × 10^−6^); here and below, correlation is provided for QuantumClone results only, although the observed trend is usually true for the three other methods.

#### Effect of the number of variants

Unexpectedly, increasing the number of variants slightly decreased the quality of clustering (*rho* = −0.17, *p — value* = 0.0104). This effect is due to an artefactual increase in the number of mutation clusters needed to explain larger numbers of observed cellular prevalence values. In other words, independently of the true clonal structure, more variants in the input result in the prediction of higher number of mutation clusters corresponding to clones.

#### Effect of the number of samples

To reconstruct clones existing in patients’ tumors with higher fidelity, recent studies advocate for sequencing of multiple samples obtained from the same patient, either by using different time points or different sites of the tumor (Schwarz et al., 2014). Indeed, we demonstrated that increasing the number of available samples significantly improved the quality of clonal reconstruction (*rho* = 0.40, *p* — *value* = 6.3 × 10^−6^).

#### Effect of sequencing depth

Variance in VAF estimation is expected to go down with the increase in sequencing depth, resulting in better mutation clustering. As expected, we observed a significant positive correlation between depth of sequencing and the quality of clonal reconstruction (*rho* = 0.67, *p* — *value <* 2.2 × 10^−16^).

#### Effect of the number of distinct clones present

We observe that the quality of clonal reconstruction declined with the increase in tumor heterogeneity, i.e., the number of distinct clones present in patients’ tumors (rho = −0.27, *p* — *value* = 8.2 × 10^−5^). As previously mentioned, in heterogeneous samples, QuantumClone showed better performance than the other tested methods.

### Creating a robust framework for clonal assignment of functional mutations

We proposed a novel concept of reconstruction of the clonal architecture in cancer combined with the attribution of functional mutations (potential drivers) to identified clones (Fig. 1). The approach is based on the different usage of *‘functional’* variants that potentially affect cell phenotype and *‘support’* variants that are used to define clones. *Support* variants can be either drivers or passengers; however, they should have high depth of coverage (> 50x in our implementation), have no strand bias and should not coincide with annotated single-nucleotide polymorphisms (SNPs). As we showed in the simulation studies (Fig. 2B) only a limited number of support variants are needed for an accurate clonal reconstruction. Therefore, in most cases, we can even afford to limit the set of support variants to those falling in regions of genotype A and AB; in such regions VAF values directly determine values of mutation cellular prevalence (Methods). *Support* variants, because they have a lower variance of observed VAF compared with other variants, are applied to define clones, i.e., *support* variants serve as input to QuantumClone or an alternative method. *Functional* mutations are defined as variants with deleterious properties that affect either genes reported in the Cancer Census List (Futreal et al., 2004) or genes from gene modules/signaling pathways that are recurrently affected by mutations in a given cancer type (Methods). At the last step of our framework, functional mutations are mapped to the clonal structure inferred from support variants based on the likelihood values.

The QuantumClone R package includes functions for both clonal reconstruction using *support* variants and assignment of *functional* mutations to the defined clones.

We propose the following options to be used in the analysis framework. Deleterious mutations can be determined using SIFT (Ng and Henikoff, 2003), PolyPhen-2 (Adzhubei et al., 2013) and FunSeq2 (Khurana et al., 2013). Gene module enrichment analysis may be performed using the R package ACSNMineR (Deveau P., Barillot E., Boeva V., Zinovyev A., Bonnet E., *In Press*) using maps and modules of the Atlas of Cancer Signalling Networks (ACSN) (Kuperstein et al., 2015) completed with the user-defined/cancer-specific modules. For the specific case of neuroblastoma, we created a ‘Neuritogenesis’ map extracted from Molenaar et al. (2012).

### Characterization of neuroblastoma clonal evolution from diagnosis to relapse: application of the QuantumClone-based framework

We applied our framework to investigate the clonal composition of neuroblastoma primary and relapse tumors and study its clonal evolution. We characterized clonal structure of tumors of 22 neuroblastoma patients (clinical data available in Suppl. Table 1). We performed WGS of constitutive DNA, diagnosis and relapse tumor samples of each patient with average depth of coverage ~ 100 x. Datasets for 15 patients out of 22 came from a previously published study (Eleveld et al., 2015). Sequencing was carried out using both Illumina HiSeq 2500 and Complete Genomics platforms. Reads were mapped to the reference hg19 genome using BWA-aln (Li and Durbin, 2009) (Illumina reads) and the internal Complete Genomics mapping tool (Complete Genomics reads). Variant calling was performed using Varscan2 version 2.3.6 (Koboldt et al., 2013).

The level of contamination by normal cells varied from 0% to 90%, and only data from 19 patients with a contamination level lower than 70% were kept for further analysis (Suppl. Table 1).

### Application of filters unifies variant call numbers across different sequencing platforms

In order to remove false positive variant calls, we used a set of stringent filters (Fig. 1, Methods). The initial number of variants in the Varscan2 output was highly dependent on the sequencing technology and platform (Suppl. Fig. 1 and 2). The number of variants called for samples sequenced by the Beijing Genomics Institute (BGI) sequencing platform was an order of magnitude higher than the number of mutations called for samples processed by the Centre National de Génotypage (CNG). Application of a set of filters based on read depth of coverage, read mappability, annotated repetitive regions (listed in Fig. 1 and Methods) allowed us to get comparable numbers of variants for further analysis. In the final list, number of mutations per sample correlated with the age of the patient (Fig. 4, Spearman’s rho = 0.54, *p — value* = 5.3 × 10^−4^; datasets tested include information from all samples kept after the evaluation of normal contamination). The stability of final numbers of predicted variants across sequencing platforms and correlation of these numbers with the age of patients validated our filtering approach. In fact, the presence of such correlation has previously been shown in neuroblastoma (Molenaar et al. 2012). In all but one neuroblastoma patient, there were more variants detected in the relapse sample than in the diagnosis sample with an average two-fold increase.

**Figure 4:**
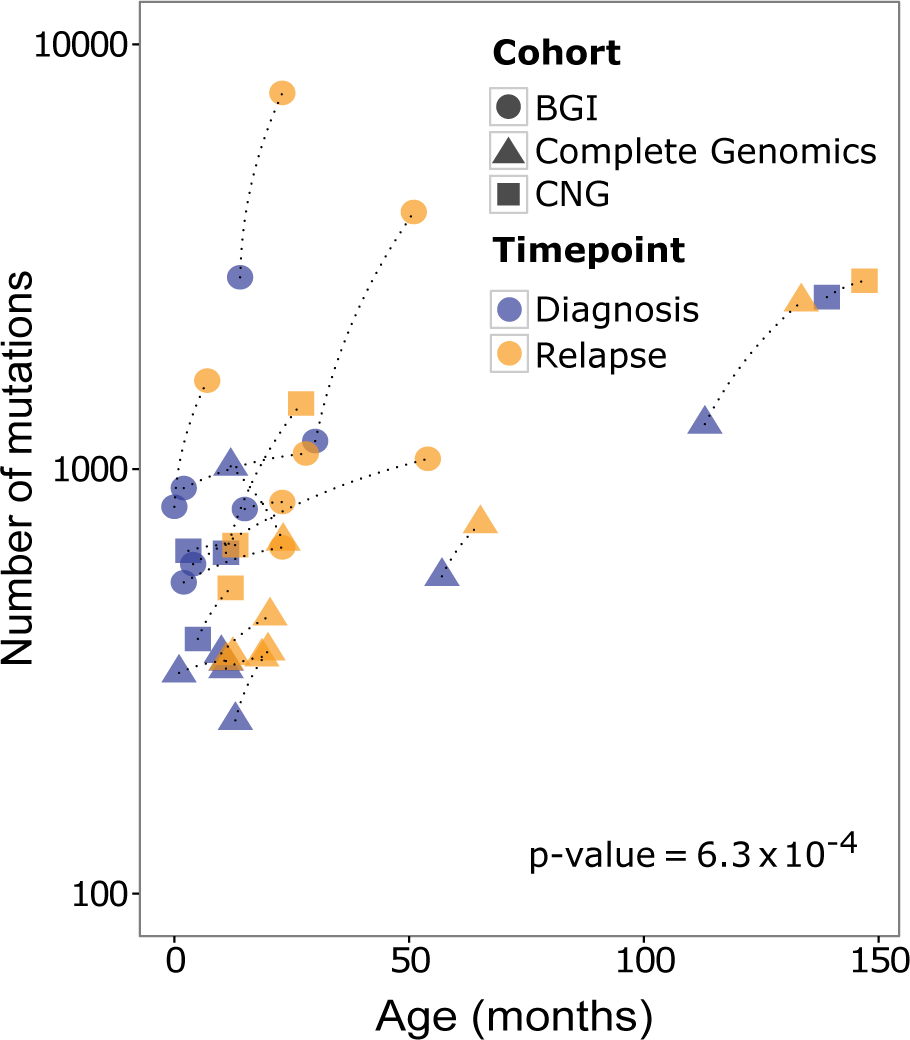
Statistics on numbers of somatic mutations called using stringent filters for diagnosis and relapse samples from 19 neuroblastoma patients. The number of somatic mutations is correlated with the age of the patient at the time of the biopsy (Spearman correlation test *p — value* = 6.3 × 10^−4^). Final mutation numbers after the filtering step do not depend on the sequencing center or sequencing technology used. Diagnosis and relapse samples from the same patient are connected by a dotted line.

### Clonal reconstruction

We applied QuantumClone on *support* variants we defined using stringent filters (Fig. 1. Step 2: Fig. 5). Across our cohort, we observed a significant association between the predicted number of clones and the number of mutations per patient (Spearman’s *rho* = 0.42, *p — value* = 0.011). However, for each given patient, the number of clones at relapse was similar to that at diagnosis, even despite the fact that the relapse samples had about twice as many mutations as the diagnosis samples (number of mutation clusters varied from zero to four with a median of three for both time points).

**Figure 5:**
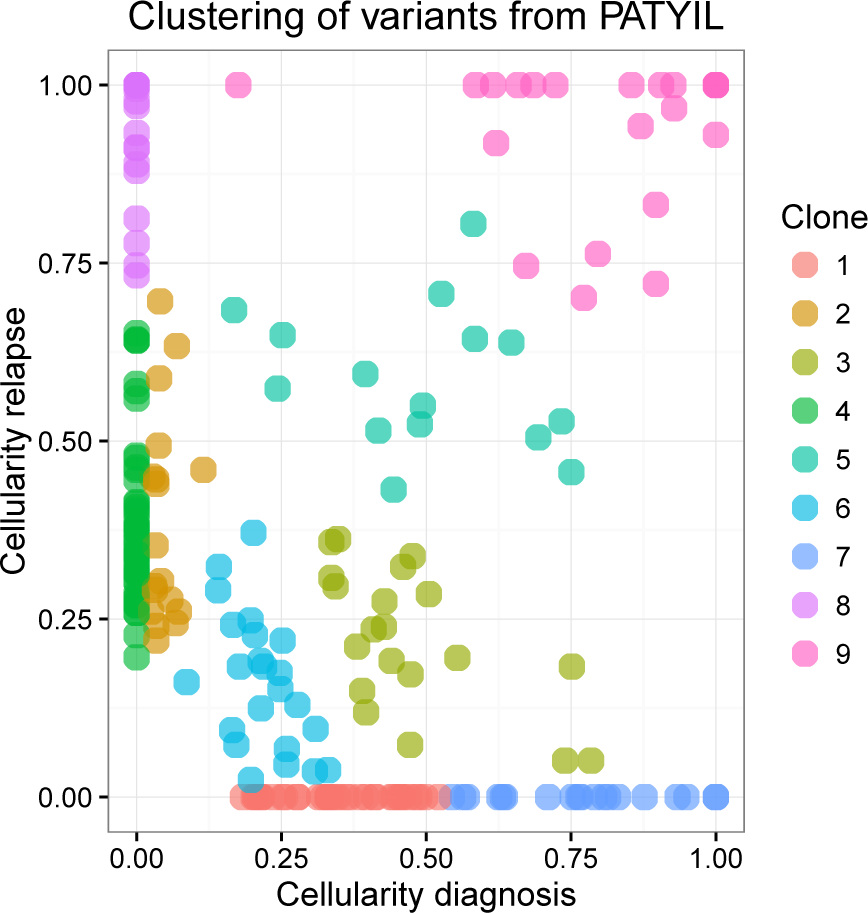
Visualization of the QuantumClone output on the PATYIL neuroblastoma patient. Nine mutation clusters corresponding to inferred clones are shown in different colors. The cellular prevalence of each mutation was assessed by QuantumClone by applying the EM algorithm on VAFs using known levels of normal contamination and locus genotype information.

We identified mutations coming from the ancestral clone (Figure 6A), i.e. the clone that gave rise to all cells in both diagnosis and relapse samples, in 84% of reconstructed clonal structures (16 out of 19 patients). The total number of *support* variants attributed to ancestral clones ranged between 3 and 120. with a median of 22.

**Figure 6:**
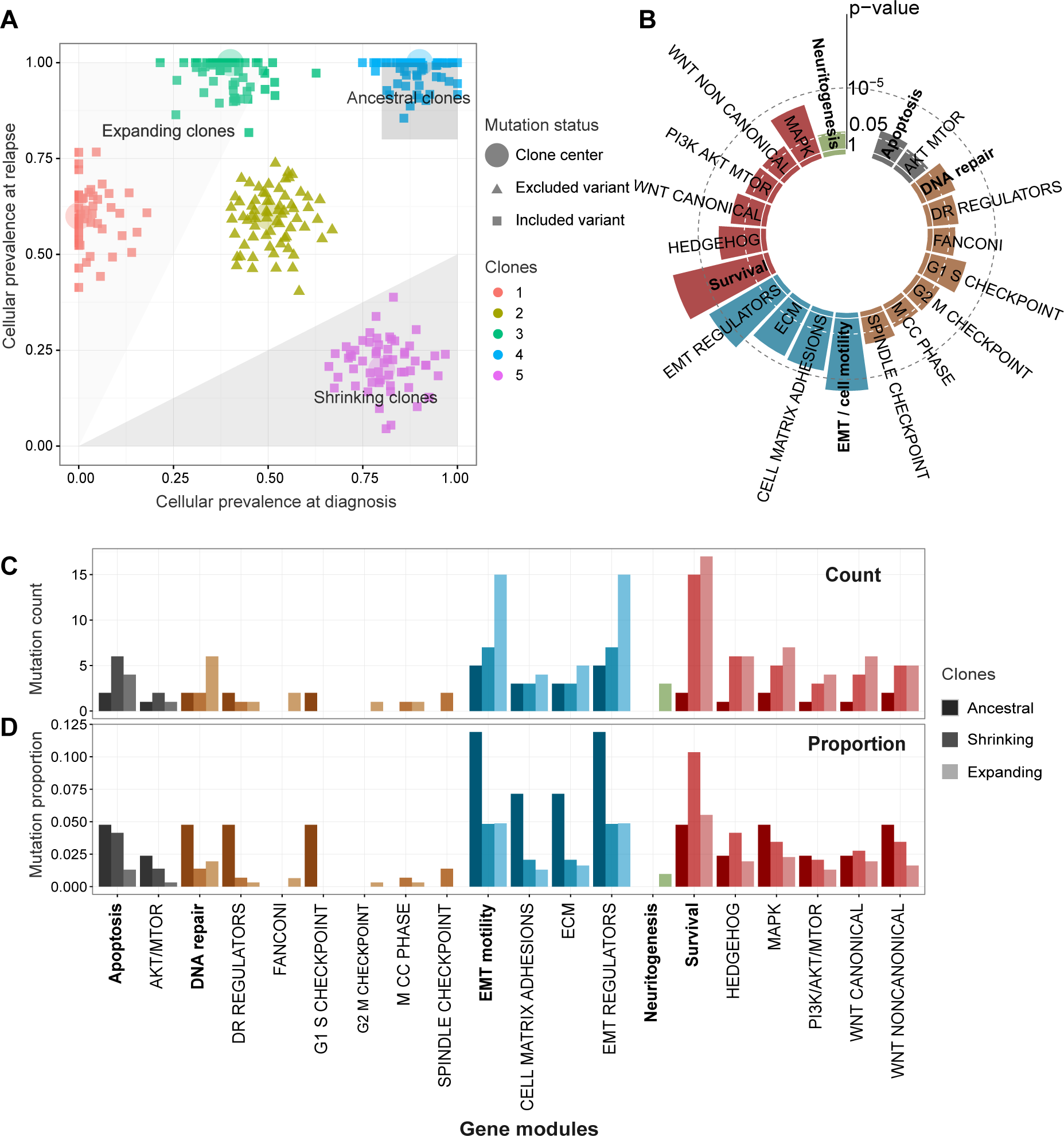
Annotation of clones in neuroblastoma and pathway enrichment analysis. **(A)** Illustration of the rule for assignment of mutations to (i) the ancestral clone (cellular prevalence of the mutation cluster exceeds 80% both at diagnosis and relapse), (ii) clones expanding after the treatment (cellular prevalence of the mutation cluster increases at least two-fold at relapse) and (iii) shrinking clones (cellular prevalence of such mutation clusters decreases at least two-fold). **(B)** Pathways enriched in deleterious mutations in neuroblastoma. General ACSN maps and the manually added neuritogenesis pathway are shown in bold: map sub-modules are shown in the color of the corresponding map. The “Cell Cycle” ACSN map does not show enrichment in mutations and is omitted from the graph. **(C)** Absolute numbers of deleterious mutations in gene maps and modules in the ancestral clones, and clones expanding and shrinking at relapse. **(D)** Proportions of *functional* mutations in each module over the total number of detected deleterious mutations in the ancestral, expanding and shrinking clones.

### Annotation of functional mutations in each sample based on the global pathway enrichment analysis

In our framework, we assumed that *functional* mutations in a given cancer type. i.e. putative drivers, should target specific signaling pathways or pathway modules (Fig. 1. Step 4). These modules can be identified as those enriched in coding deleterious mutations across the cohort. Tims, we mapped the total of 541 deleterious mutations obtained with tolerant filters (Fig. 1. Methods) to the ACSN maps and detected recurrently altered gene modules using the ACSNmineR package. Overall, five general gene maps (apoptosis. DNA repair. EMT/cell motility, cell survival and neuritogenesis) and their 15 gene modules were found to be enriched in mutations (threshold 0.05 on the p-value corrected to account for multiple testing with the Benjamini-Hochberg False Discovery Rate correction, corresponding to the q-value)) (Figure 6B. Suppl. Table 2). Next, deleterious mutations were annotated as *functional* when corresponding genes were included in the enriched pathways, or when such genes belonged to the Cancer Census list. The resulting number of *functional* mutations per sample varied from 1 to 23.

At this step, among the cell survival modules, the highest enrichment in putative driver mutations was observed for the MAPK pathway (*q — value ≤* 7.2 × 10^−5^). In addition, we detected significant enrichment in *functional* mutations of both canonical and non-canonical WNT pathways (*q* — *value ≤* 3.8 × 10^−3^ and ≤ 1.49 × 10^−2^, respectively), and of the PI3K/AKT/mTOR and Hedgehog gene modules (*q* — *value ≤* 1.3 × 10^−2^ and ≤ 4.5 × 10^−4^, respectively). As for the modules of other maps, genes coding for the EMT regulators were also significantly affected by the deleterious mutations in our cohort of relapsed neuroblastoma patients (*q* — *value* ≤ 1.2 × 10^−6^).

As a control, we ran ACSNMineR on synonymous variants passing the same filters as the non-synonymous ones (1060 synonymous mutations in 771 genes). Enrichment analysis provided no modules enriched in synonymous variants (*minimal p—value* > 0.28), thus confirming the biological significance of the discovered enrichment in functional mutations for 20 ACSN modules and maps. Strikingly, similar enrichment analysis of intronic variants showed recurrently affected pathways similar to those affected by coding deleterious mutations, such as EMT-motility (*Odds ratio* = 5.3, *p — value <* 10^−20^) or cell survival (*Odds ratio* = 4.0, *p — value <* 10^−20^), highlighting the possible role of intronic SNVs in neuroblastoma tumorigenesis.

### Assignment of *functional* mutations to the identified clonal structure

Using the results of the mapping of *functional* mutations on the clonal structure detected for each patient by QuantumClone (Fig. 1, Step 5), we annotated mutations as (*i*) those belonging to expanding clones, (*ii*) those belonging to shrinking clones, and (*iii*) those belonging to ancestral clones (Fig. 6A). Overall, 53%, 31% and 4,8% of all *functional* mutations fell in these three categories.

For the majority of samples (16 out of 19 patients), we could not detect *functional* mutations in the ancestral clone. But, interestingly, samples with identified ancestral *functional* mutations contained variants in the *MYC* and *AKT2* genes, which are known to act as drivers in many cancers. Moreover, the ancestral clones detected in our samples, often contained a higher proportion of putative driver mutations affecting the DNA repair, EMT/cell motility and other pathways than any other clones including those expanding at relapse (Fig. 6B and 6C).

### Analysis of pathways enriched in *functional* mutations in shrinking and expanding clones

Assignment of mutations to clones shrinking or expanding after the treatment resulted in the identification of 33 and 56 possible driver mutations in these clone types, respectively. Expanding clones had more deleterious mutations targeting genes from four general maps (DNA repair, EMT/cell motility, cell survival and neuritogenesis) (Fig. 6C). Similarly, in these clones, most of the corresponding gene modules (e.g., MAPK, WNT canonical or PI3K/AKT/mTOR) were also more frequently targeted. Although the absolute number of *functional* mutations in the expanding clones was about twice as high as in the shrinking ones, the proportion of such mutations among all deleterious ones was not significantly different between the shrinking and expanding clones (Fig. 6D).

In addition to the MAPK pathway mutations previously reported for the relapse neuroblastoma samples (e.g., mutations in the *NRAS* gene (Eleveld et al., 2015)), in the expanding clones we observed coding variants affecting additional MAPK pathway genes such as *NFE2L1*, *PIM1*, *FLT4* and *RSK*. These genes have been shown to be involved in tumorigenesis of prostate (Yu et al., 2015), colorectal (Xiao et al., 2014), breast (Malinen et al., 2013) and other cancers.

## Discussion

Here we propose a pathway-based framework to detect functional mutations in cancer samples and associate the mutations to their corresponding clonal structure. The central part of our framework is represented by the QuantumClone method, which allows reconstruction of clonal populations based on both variant allele frequencies and genotype information. QuantumClone showed stable results on simulated data significantly outperforming other methods in difficult settings such as highly contaminated samples, heterogeneous tumors and relatively low depth of sequencing coverage.

Our analysis framework is based on two central ideas. First, high-reliability passenger mutations must be used to reconstruct the clonal structure of tumors samples; then, low coverage functional mutations (with high variance in VAFs) should be mapped onto the inferred clonal structure. Second, we suggest to limit the set of functional mutations to those in genes known to be associated with cancer (e.g., Cancer Census genes) or to those in genes from gene modules/pathways frequently disrupted in a given cancer type (Fig. 1).

We apply the proposed analysis framework to decipher clonal structure in neuroblastoma and assign to clones possible driver mutations. We detect 15 pathways as being altered by mutations in neuroblastoma. We identify genes associated with DNA repair, cell motility, apoptosis and survival to be enriched in functional mutations in neuroblastoma.For relapsed neuroblastoma samples, we recover the previously reported enrichment of mutations in the MAPK signaling pathway (Eleveld et al., 2015), while complementing this knowledge with discovery of accumulation of functional mutations at the relapse in such functional gene modules as PI3K/AKT/mTOR, WNT, Hedgehog signaling and modules consisting of genes responsible for cell-matrix adhesion and epithelial-mesenchymal transition (EMT).

We highlight the lack of enrichment in mutations of the cell cycle pathway in this pediatric cancer. This could be explained by the biological context of neuroblastoma that may already favor proliferation, which accompanies organism development, thus limiting the necessity to disrupt cell cycle mechanisms. However, this observation should be in the future confirmed on a larger cohort.

For the majority of neuroblastoma patients, we did not identify driver mutations in the ancestral clone. This is in line with the current understanding of neuroblastoma as a type of cancer driven by copy number alterations. In fact, it has been known that the aggressiveness of neuroblastoma is highly associated with changes of the chromosomal copy number profile (MYCN amplification, lp, 3p, llq deletions, partial gain of chr17) (Janoueix-Lerosey et al., 2010). And indeed, copy number profiling often detected gain and losses in these recurrently affected chromosomal regions both in the diagnosis and relapse samples from our cohort (Suppl. Fig. 3).

For most of our samples, we did not succeed in reliably reconstructing the phylogeny of clonal evolution based on cellular prevalence values for identified mutation clusters (clones). Some contradictions between cellular prevalence values between diagnosis and relapse, as well as disappearance at relapse of many potential driver mutations seemingly present in the ancestral clone at diagnosis, may be due to tumor heterogeneity and the fact that biopsies were taken from different tumor sites. This situation has been termed “illusion of clonality” (Bruin et al., 2014). However, the fact that for most of the samples we observed a number of shared mutations between diagnosis and relapse and that copy number breakpoints were consistent between the two time points ensures that there is a common phylogeny between diagnosis and relapse in neuroblastoma (Bollet et al., 2008).

In the application of our framework to neuroblastoma sequencing data, we excluded information about SVs and indels. The reason for this was that the analysis of clonal structure is based on the number of sequencing reads supporting each genetic variant. While we suppose that the number of reads with a mismatch mutation is proportional to the number of DNA molecules harboring this variant, we expect that due to read mapping issues the fraction of reads indicating an indel or a large SV will be generally lower than the actual proportion of DNA molecules with the rearrangement. Eviction of large and small SVs seemingly resulted in a decrease in sensitivity of the detection of genetic driver events. In our neuroblastoma data, a large proportion of observed clones did not contain any predicted driver mutation. In the future, the sensitivity can be improved by using higher depth of coverage data and combining the paired-end datasets with reads produced with the mate-pair protocol or with long PacBio reads.

The proposed framework can be applied in the future to any type of cancer. The pre-requirements are sufficient number of candidate mutations (at least 50 mutations per sample) and a minimal read depth of coverage of 50 x. These requirements are usually met by WGS or whole exome sequencing datasets. Our simulation results show that increasing the number of mutations used for clonal reconstruction above 50 does not improve significantly the clonal reconstruction accuracy provided that mutations specific for every clone are present in the input.

## Methods

### Datasets

#### Patient selection and collection of tumor samples

The inclusion criteria for this study were histopathological confirmation of neuroblastoma at original diagnosis and the presence of biopsy material from a subsequent relapse specimen. Patients were included in this study after an informed consent was obtained from parents or guardians, with oversight from the ethics committees ‘Comité de Protection des Personnes Sud-Est IV’, reference (.07-95 (.12-171. and ‘Comité de Protection des Personnes Ile-de-France’, reference 0811728 in France, the review board at the Children’s Hospital of Philadelphia and review boards at other Children’s Oncology Group sites that submitted samples for patients on this study in the United States. In total we obtained material for 22 neuroblastoma patients (tumor tissue at diagnosis, relapse and constitutional DNA, Suppl. Table 1).

#### Whole-genome sequencing of neuroblastoma samples

In the framework of this study, we carried out Illumina paired-end sequencing for 7 novel neuroblastoma patients (corresponding to 21 samples). Data for 15 patients were taken and reanalyzed from the previous study (Eleveld et al., 2015). DNA from 7 patients from the previous study and 7 new ones have been sequenced using Illumina HiSeq 2500 instruments to an average depth of coverage of 80× by Beijing Genomics Institute (BGI) and the Centre National de Génotypage (CNG) respectively. For 8 patients out of 15 previously reported, whole-genome sequencing was performed by Complete Genomics with an average read depth of coverage of 50×. DNA material for each patient (lymphocytes, primary tumors and relapse tumors) was in each case sequenced using the same sequencing platform (see Suppl. Table 1 for more detail).

#### Data processing

Sequenced reads were mapped to the human genome hg19 using BWA and the internal Complete Genomics tools for Illumina and Complete Genomics datasets respectively. Reads from datasets sequenced using the Illumina platform were realigned around indels with the Genome Analysis ToolKit (GATK) (McKenna et al., 2010), followed by a base recalibration. Due to the inherent structure of Complete Genomics reads, which contain an effective deletion relative to their corresponding genomic library, the indel realignment step was skipped for the Complete Genomics samples.

### Variant calling and filtering

Mutations were called using Varscan2 (Koboldt et al., 2013). Two sets of variants were created for each patient (see Fig. 1) using tolerant and stringent filtering options. The ‘stringent’ set was further used for clonal reconstruction, while the ‘tolerant’ one was used for inference of recurrently altered pathways.

Tolerant filters for somatic mutations included those on minimal depth of coverage (30×), minimal percentage of reads supporting the mutation (10%). In addition, mutations were required to be located in regions of high local mappability (36 bp mappability), outside of repeat and duplicated genomic regions (assessed by the UCSC repeat and segmental duplication region tracks). We further filtered mutations that created a stretch of four or more identical nucleotides. Finally, we only kept mutations located in regions where the genotype evaluated by Control-FREEC was available.

To obtain a set of high condence mutations, we required the minimal depth of coverage of 50×. We filtered out variants corresponding to polymorphisms present in more than 1% of the population (snpl38, lOOOGenomes, esp6500) except if it was a known cancer related variant (COSMIC database for coding and non-coding mutations). Strand bias was also tested by the Fisher exact test (in addition to the test in Varscan2) to reduce the number of informative mutations to a maximum of 500. We restricted the analysis to AB regions in case when after such filtering we kept more than 100 mutations.

### Copy number analysis

Copy number alterations in patients were detected using the Control-FREEC method (Boeva et al., 2012) (version 7.2) (Suppl. Fig. 3). We selected the main ploidy value so that the predicted copy number and B-allele frequency profiles were consistent. Control-FREEC also provided estimations of the level of contamination by normal cells, which, after manual confirmation, was further used for the clonal reconstruction.

Three samples with the estimated proportion of contamination by normal cells higher than 70% were excluded from the further analysis (NB0784:diagnosis, NB1434:relapse and NB1471:relapse).

### Comparison of clonal reconstruction between QuantumClone and existing methods

#### Data simulation

In silico validation data were generated using the QuantumCat method from package QuantumClone (version 0.15.12.10). QuantumCat simulates genomic mutations, copy number alterations and corresponding VAFs. It relies on the following set of rules:

1. A binary phylogenetic tree is created to simulate the clonal architecture of the tumor. The mutation cellular prevalence values correspond to the nodes and leaves of the phylogenetic tree.
2. Cellular prevalence values of mutations from each clone are independent across tumor samples. However, the cellular prevalence of each clone should be always coherent with the phylogenetic tree.
3. The allelic copy number of all mutation loci was set to AB in the tests carried out to compare QuantumClone, sciClone and pyClone (Figure 2B). For QuantumClone validation on triploid and tetraploid genomes (Figure 3), the number of chromosomal copies bearing each mutation was randomly assigned between one and the number of A-alleles for the locus considered. Generation of the genotype, number of chromosomal copies, normal contamination and cellular prevalence of a mutation allows for the computation of the exact VAF, which is the cellular prevalence (taking into account the contamination by normal cells) times the number of copies of the mutations divided by the number of copies of the locus in each cell. On the other hand, the observed VAF is determined by the ratio of the number of reads supporting the mutations divided by the read depth of coverage. Local depth of coverage at each given position was generated by the negative binomial distribution centered on the target depth of sequencing, fitted on experimental data. The number of reads supporting a mutation was simulated from the binomial distribution with the probability of success equal to the exact VAF.

#### Program, versions and parameters

We used SciClone version 1.0.7 with the following changes to the default parameters: maximal number of clusters was set to 10 and the minimal depth of coverage was set to 0.

PyClone version 0.12.9 was used with the following parameters : 10 iterations of the Markov chain Monte Carlo, alpha and beta parameters in the Beta base measure for Dirichlet Process set to 1, concentration prior shape set to 1 and the rate parameter in the Gamma prior on the concentration parameter set to 0.001. We used the default Beta binomial distribution with precision parameter set to 1000, prior shape set to 1, rate to 0.0001, and proposal precision set to 0.01.

We used an implementation of the *k*-medoids algorithm provided by the R package “fpc”, version 2.1.10, with a range of clusters between 2 and 10.

For clonal reconstruction, QuantumClone version 0.15.12.10 was used with default parameters except for the the maximal number of clusters, which was set to 10. For clonal reconstruction of neuroblastoma data, mutations used to compute centers of clusters (corresponding to clones) were selected using the stringent set of filters. Copy number information from Control-FREEC (version 7.2) was also passed to the algorithm as well as the predicted value of contamination by normal cells.

For simulated data, quality of clustering was assessed by using Normalized Mutual Information (NMI), which is given for a group of clones Ω and a group of reconstructed clusters &#x2102;:

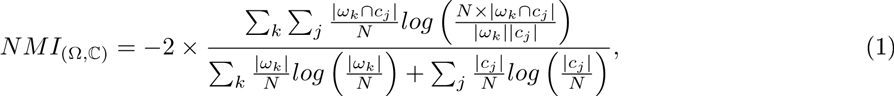

where *N* is the number of mutations observed, |*ω_k_* | the number of mutations in clone *k*, and |*c_j_* | the number of mutations attributed to cluster *j*.

### Clonal reconstruction

In this section we describe QuantumClone, a method we have developed for the clonal reconstruction of a tumor. QuantumClone performs clustering of cellular prevalence values of mutations defined by:

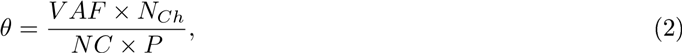

where *θ* is the cellular prevalence, *N_Ch_* the number of copies of the corresponding locus, *NC* the number of chromosomal copies bearing the mutation, and *P* the tumor purity. For instance, only in case of a purely diploid tumor without loss of heterozygosity (LOH) regions, with no contamination of the sample by normal cells, the cellular prevalence is equal to 2 × *VAF.* The latter assumption has been frequently used in cancer studies (Schramm et al., 2015). As we do not have information about the number of chromosomal copies bearing a mutation, our approach was to compute each possible value of cellular prevalence associated with the mutation VAF. For example, a mutation can have a VAF of 1/3 in a locus of genotype AAB when it is present in 100% of tumor cells on a single chromosome copy and when it is present in 50% of tumor cells on two chromosomes. Yet the latter case is rather improbable. Each mutation thus corresponds to several possible values of cellular prevalence; each solution is associated with a value of *NC*. In order to address the problem of non-uniqueness of solution, we use an EM algorithm based on the probability to observe a specific number of reads confirming a mutation given the number of reads overlapping the position, the contamination and the cellularity of a clone. In more detail, we attribute to each possibility a probability to observe *f* reads supporting the variant given that the latter belongs to a clone of cellular prevalence *θ*, based on a binomial distribution:

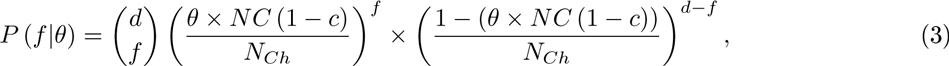

where

- *d* is the depth of coverage of the variation
- *f* is the number of reads supporting the variant
- *c* is the sample contamination by normal cells

We can then write the log likelihood function to maximize:

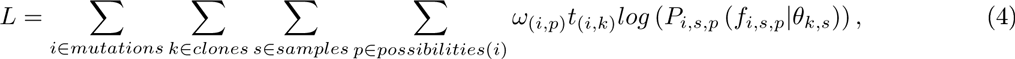

where *ω*_*i*,*p*_ are weights of the possibility computed for a corresponding genotype *xAyB* (major allele *A* is present *x* times and the minor allele *B* is present *y* times):

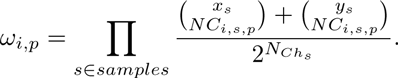

By adding weights that for each variant sum to one, we favour mutations with the lowest number of copies. Each mutation is then attributed to its most likely possibility, which is the possibility with highest probability to belong to a clone. In the situation described above (a variant in a AAB region with the VAF of 1/3), this approach would assign probabilities of 2/3 and 1/2 to the presence of the mutation in 100% and 50% of cells respectively. However, if there is a second mutation present, for example, in a locus of genotype AB with a VAF of 1/2 and thus having unambiguously cellular prevalence of 100%, the rst mutation will have a high density of probability for a cellular prevalence of 100% and our approach will assign both mutations to the same cluster corresponding to the same cellular prevalence (100%).

The number of clones is determined by minimization of the Bayesian Information Criterion (BIC). Priors can be provided by the user, randomly generated or determined by the *k*-medoids clustering on mutations in A and AB sites when the latter contain enough mutations.

#### QuantumClone-Single and QuantumClone-Alpha variants of QuantumClone

To test accuracy of predictions for the number of copies with a variant and selection of the most likely mutation cellular prevalence, we designed two alternatives of the QuantumClone method: QuantumClone-Single and QuantumClone-Alpha. QuantumClone-Single is a modification of QuantumClone that assigns to all variants a single copy state. QuantumClone-Alpha uses the same EM algorithm as the default version of QuantumClone, except for the selection of the most likely mutation where it chooses the possibility maximizing the quantity *q*:

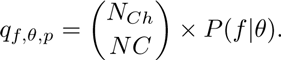

### Analysis of mutation enrichment in signaling pathways and gene modules

ACSNMineR (Deveau P., Barillot E., Boeva V., Zinovyev A., Bonnet E., *In Press*) version 0.16.01.29 was used to compute gene modules and pathways enriched in deleterious mutations. Gene modules included by default in ACSNMineR come from the manually curated Atlas of Cancer Signalling Networks (ACSN) (Kuperstein et al., 2015). In addition to the ACSN modules, we calculated mutation enrichment in a set of neuritogenesis genes frequently mutated in neuroblastoma (Molenaar et al. (2012), Suppl. Table 8). We called deleterious mutations as stop-gain mutation or variants that were predicted as possibly damaging or deleterious by SIFT (Ng and Henikoff, 2003), PolyPhen-2 (Adzhubei et al., 2013), or FunSeq2 (Khurana et al., 2013).

To get a list of genes to use as an input to ACSNMineR, we pooled mutations from all neuroblastoma patients; genes mutated at least once were included in the final list. Modules with a p-value lower than 0.05 after Benjamini-Hochberg correction were considered as enriched.

## Data access

The whole-genome sequencing data have been deposited at the European Genome-phenome Archive (EGA) under accession number EGAS00001001184 for the French cases sequenced at BGI and under accession number EGAS00001001825 for the French cases sequenced at CNG. Sequence data for the US cases are available in the database of Genotypes and Phenotypes (dbGaP) under accession number phs000467.

QuantumClone is available at https://github.com/DeveauP/QuantumClone/ and can be downloaded as an R package from the CRAN repository.

## Acknowledgments

GS and her team were supported by the Annenberg Foundation and the Nelia and Amadeo Barletta Foundation. Funding was also obtained from SiRIC/INCa (Grant INCa-DGOS-4654) and from the CEST of Institut Curie. This study was also funded by the Associations Enfants et Santé, Association Hubert Gouin Enfance et Cancer, Les Bagouz à Manon, Les amis de Claire. VB and her team were supported by the ATIP-Avenir Program, the ARC Foundation and the “Who Am I?” Project. EB was supported by the ABS4NGS project of the French Program ‘Investissement d’Avenir’. Sequencing of French samples was carried out in a collaboration of Institut Curie with CEA/IG/CNG financed by France Génomique infrastructure, as part of the program “Investissements d’Avenir” from the Agence Nationale pour la Recherche (contract ANR-10-INBS-09). JM and his team were supported in part by US National Institutes of Health grants RC1MD004418 to the TARGET consortium, and CA98543 and CA180899 to the Children’s Oncology Group. In addition, this project was funded in part with Federal funds from the National Cancer Institute, National Institutes of Health, under Contract No. HHSN261200800001E. The content of this publication does not necessarily reflect the views of policies of the Department of Health and Human Services, nor does mention of trade names, commercial products, or organizations imply endorsement by the U.S. Government.

## DISCLOSURE DECLARATION

We have no conflict of interest to declare.

